# PACAP Signaling Network in the Nucleus Accumbens Core Regulates Reinstatement Behavior in Rat

**DOI:** 10.1101/2025.03.17.643720

**Authors:** Samrat Bose, Gregory Simandl, Evan M. Hess, Linghai Kong, Nicholas J. Raddatz, Brian Maunze, SuJean Choi, David A. Baker

**Author notes:** Co-first authors due to equal contributions (Samrat Bose, Gregory Simandl, Evan M. Hess, Linghai Kong).

## Abstract

Cocaine use disorder (CUD) lacks FDA-approved treatments, partly due to the difficulty of creating therapeutics that target behavior-related neural circuits without disrupting signaling throughout the brain. Recent evidence highlights the therapeutic potential of targeting gut-brain axis components, such as GLP-1 receptors, to modulate neural circuits with minimal central nervous system disruption. Like GLP-1, pituitary adenylate cyclase polypeptide (PACAP) is a component of the gut-brain axis that regulates behavior through a network spanning the gut and brain. Here, we investigated the potential existence and function of an endogenous PACAP signaling network within the nucleus accumbens core (NAcc), which is a structure that integrates emotional, cognitive, and reward processes underlying behavior. We found that PACAP and its receptor, PAC1R, are endogenously expressed in the rat NAcc and that PACAP mRNA is present in medial prefrontal cortical projections to the NAcc. Behaviorally, intra-NAcc infusions of PACAP (100 pm) did not induce seeking behavior but blocked cocaine-primed reinstatement (10 mg/kg, IP). Intra-NAcc PACAP also inhibited reinstatement driven by co-infusion of the D1 receptor agonist (SKF 81297, 3 µg) but not the D2 receptor agonist (sumanirole, 10 ng). These findings are significant since D1 and D2 receptor activities in the NAcc govern distinct behavioral mechanisms indicating precise actions of PACAP even within the NAcc. Future research should examine whether NAcc PACAP signaling can be selectively engaged by peripheral gut-brain axis mechanisms, potentially unveiling novel therapeutic approaches for CUD and related disorders.

## Introduction

Impaired behavioral control underlies chronic disorders such as addiction, stress-related conditions, and other psychiatric diseases, posing a significant public health challenge^1,2^. Despite decades of research linking impaired behavioral control to dysfunction in neural circuits that integrate emotional, cognitive, and reward processes^3–9^, current treatment options remain limited. For example, cocaine use disorder (CUD) lacks a single FDA-approved medication.

Developing therapies for central nervous system (CNS) disorders is far more challenging than for chronic diseases affecting other tissues^10,11^. A key barrier to CNS therapeutic development is that many conventional experimental therapeutics act at neurotransmitter receptors, transporters, or related mechanisms that are expressed by circuits distributed throughout the brain^12^. Hence, brain-wide distribution of experimental therapeutics targeting conventional neurotransmitter systems will impact the activity of circuits that produce therapeutic outcomes as well as those that produce onsite or offsite adverse effects. Ultimately, the development of safe and effective treatments may require identifying novel strategies that effectively modulate behavioral control circuits selectively while avoiding broad CNS disruption.

The recent clinical success of peripherally restricted GLP-1 receptor agonists underscores the therapeutic potential of gut-brain axis mechanisms to regulate brain circuits^13,14^. GLP-1 signaling selectively modulates feeding-related circuits via relays, including vagal afferents and circumventricular structures, to influence central neural networks^13–19^. As a result, peripherally restricted GLP-1 agonists can effectively treat obesity, type 2 diabetes, and potentially additional indications while producing minimal CNS side effects^20–23^. However, fully realizing the therapeutic potential of the gut-brain axis in CNS disorders requires a deeper understanding of both targetable peripheral mechanisms and modifiable CNS processes.

Pituitary adenylate cyclase-activating polypeptide (PACAP) and GLP-1 are both in the same family of structurally-related peptides that are distributed peripherally and centrally^24,25^. However, PACAP exhibits a much broader distribution in the central nervous system as it is expressed in cortical, subcortical, and midbrain circuits. Hence, central PACAP may be a key central component of the gut brain axis given the complex circuitry underlying CUD and related conditions^26–31^. PACAP has been implicated in stress-related relapse behaviors and alcohol consumption in specific rodent strains^32–34^, but its role in cocaine-seeking and, more specifically, in the NAcc remains unclear.

Given the NAcc’s role in integrating cognitive, emotional, and reward processes^3,35–39^, we hypothesized that PACAP acts endogenously within the NAcc to regulate cocaine-seeking behavior. This is an important area of research since the NAcc integrates cortical, subcortical, and brainstem circuits that underlie the cognitive, emotional, reward processes involved in motivated behavior. Here, we investigated whether an intrinsic PACAP signaling network exists in the NAcc and whether its activation modulates reinstatement of cocaine-seeking behavior.

## Materials and Methods

### Animal care and usage

Male Sprague Dawley rats (70-150 days) were individually housed in a temperature and humidity-controlled room, maintained on a 12:12 light/dark cycle with lights-off at 0700. Rats had ad libitum access to food and water, except during the self-administration protocol. All procedures and protocols were approved by the Institutional Animal Care and Use Committee at Marquette University and adhered to the guidelines set forth by the National Institutes of Health.

### Western Blotting

Rats were decapitated, and 1mm^3^ NAcc tissue punches were lysed in 200 μl of ice-cold lysis buffer (RIPA buffer with EDTA, Halt protease, and phosphatase inhibitor cocktail, Pierce; Rockford, IL, USA). Protein extraction involved syringe homogenization (26-gauge needle, 10 passes) and sonication (three pulses, 50% power) on ice. Homogenates were centrifuged (10,000×g, 10 min, 4°C), and supernatants were collected. Protein levels were determined using a bicinchoninic acid (BCA) assay (Cat#23252, Thermo Scientific, IL, USA). A total of 30 μg protein was separated on 8% SDS-PAGE and transferred to a polyvinylidene fluoride membrane. Membranes were blocked (5% BSA, 0.1% Tween-20) and incubated overnight (4°C) with anti-PAC1R antibody (Cat#AVR-003, Alomone Labs, Jerusalem, Israel), followed by secondary antibody incubation (1 hour, room temperature; Cat#7074, Cell Signaling, MA, USA). Bands were visualized using the Odyssey Fc Imaging System. Specificity was verified by re-probing blots with and without a synthetic PAC1R peptide (Cat#BLP-VR003, Alomone Labs, Jerusalem, Israel).

### Microdialysis and PACAP estimation

A guide cannula (22g, Protech International, Roanoke, VA, USA) was implanted in the NAcc (AP +1.2mm, ML ±2.4mm, DV −6.7mm). Peptide-recovery microdialysis probes (Eicom Atmos system, San Diego, CA, USA) with pressure-canceling ports and high-molecular-weight permeable membranes were inserted into the cannulas and perfused with dialysis buffer (1 µL/min) for 2 hours. The probes were perfused with a dialysis buffer (153.5 mM NaCl, 4.3 mM KCl, 0.71 mM CaCl₂, 0.41 mM MgCl₂, 1.25 mM glucose, 0.15% BSA) at 1 µL/min and stabilized for 2 hours. Samples were then collected every 20 minutes for 2 hours, pooled into a preservative solution (PBS, 0.5% Tween-20, 2× Halt Protease and Phosphatase Inhibitor Cocktail; ThermoScientific, Waltham, MA, USA), and analyzed by ELISA (#MBS2516345, MyBioSource, CA, USA). Unstimulated extracellular PACAP levels ranged from 83–267 pM. Adjusting for probe recovery (<10%), basal levels are estimated to be in the low nM range.

### CTb retrotracer injection

Fluorescently conjugated cholera toxin subunit B (CTb 594, 1% dissolved in 1X Phosphate Buffer; Invitrogen, CA, USA) was used to trace afferent projections to the NAcc core. Using standard stereotaxic surgical procedure 300 nL of CTb was injected into the NAcc (AP +1.2 mm, ML ±2.4 mm, DV −6.7 mm, at 7 °) at a rate of 50 nL/min. Animals recovered for at least 1 week prior to their inclusion in experiments.

### In Situ Hybridization

Rats injected with CTb in the NAcc core were euthanized by rapid decapitation and brains were immediately frozen on dry ice. Brains were coronally sectioned on a cryostat at 12 μm thickness, mounted onto electrostatically clean slides, and stored at −80°C. Before hybridization, sections were postfixed in 4% paraformaldehyde, rinsed in 0.1 M PBS (pH 7.4), equilibrated in 0.1 M triethanolamine (pH 8.0), and acetylated in triethanolamine containing 0.25% acetic anhydride. Sense and antisense riboprobes targeting PACAP transcripts were generated via in vitro transcription, diluted in a hybridization cocktail (Amresco, Solon, OH, USA) with tRNA, and hybridized overnight at 55°C with FITC-labeled riboprobes. Slides were then treated with RNase A, washed in 0.1× SSC at 65°C for 30 minutes, and incubated overnight at 4°C with an antibody against FITC conjugated to horseradish peroxidase (HRP; Roche, Indianapolis, IN, USA). Riboprobe signals were amplified using the TSA-Plus fluorophore system with fluorescein (PerkinElmer; Waltham, MA, USA). Images were acquired via confocal microscopy (Nikon Instruments Inc., Melville, NY, USA) and analyzed with NiS Elements software. FIJI (ImageJ 2) was used to reassign red fluorescence to magenta and overlay confocal images onto a standard rat brain atlas via the Big Warp Viewer plugin to visualize CTb spread and PACAP mRNA co-expression.

### Canula implantation for drug delivery to the nucleus accumbens

Bilateral guide cannulas targeting the NAcc core were implanted surgically in the brain under anesthesia with stereotaxic coordinates (AP +1.2 mm, ML ±2.4 mm, DV −4.7 mm, at 7 °). Cannulas were secured in position on the skull with screws and dental cement, with injectors extending 2 mm beyond cannula tips.

### Jugular catheter surgery

Rats underwent surgical implantation of chronic indwelling jugular catheters terminating in the right atrium for cocaine self-administration under isoflurane anesthesia (inducted by 5% isoflurane and maintained with 2.5%). A silicon-tubing catheter (0.31 mm inner diameter, 0.64 mm outer diameter) was inserted into the right superior vena cava and sutured to the vein. Custom indwelling catheters were constructed of rounded tip polyurethane tubing (Access Technologies, Skokie, IL, USA) attached to a 22-gauge back mount pedestal with attached mesh (Protech International, Roanoke, VA, USA). The exit port was positioned 2 cm posterior to the scapulae. Post-surgical care included analgesia (meloxicam, 1 mg/kg, s.c.) and antibiotics (cefazolin, 100 mg/kg, i.v.) for pain and infection prevention, respectively. Catheters were flushed daily through the experiment duration with bacteriostatic heparinized saline to maintain patency and capped with Tygon tubing each time the leash/delivery line assembly was disconnected. Rats were allowed a seven-day recovery period before the start of experimental procedures^40,41^.

### Cocaine Self-Administration Training

Rats were trained on a fixed ratio 1 (FR1) reinforcement schedule for daily 2-hour self-administration sessions. Pressing the active lever extinguished the house light, activated a cue light, and delivered either a sucrose pellet (food training) or a cocaine infusion (0.5 mg/kg/inf, i.v.). Water was available ad libitum, but food was restricted to 20 g rat chow post-session. Initially, rats were trained to press for sucrose until achieving >50 presses with <15% variation over three consecutive sessions. During acquisition, training continued until rats received at least 12 daily infusions with <15% variation across three sessions. Rats then progressed to maintenance under long-access conditions (1.0 mg/kg/infusion; 6 hours/day for 12 sessions), followed by a 7–10 day withdrawal period during which bilateral guide cannulas were implanted, and rats were allowed to recover before extinction training. During extinction, pressing the active lever activated the cue light and syringe pump but delivered no infusion. Extinction criteria were set at two consecutive days of ≤15 lever presses, followed by a mock injection. If the mock injection induced >15 responses, extinction criteria were reset, and an additional mock injection was performed to confirm stability.

### Reinstatement testing

Rats received 10 mg/kg cocaine (intraperitoneal; IP) immediately prior to testing. Pharmacological interventions included bilateral intracranial infusions of vehicle, PACAP 1-38, SKF 81297, or sumanirole maleate at 0.125 µl/min (total volume: 0.5 µl) 10 minutes before testing, with lever pressing recorded for 2 hours.

### Histology

At the completion of self-administration experiments, rats were perfused with saline followed by 4% paraformaldehyde for fixation. The brains were sectioned into 150 μm slices using a Vibratome, stained with cresyl violet, and imaged. Brain images were aligned to a rat brain atlas (Paxinos and Watson, 6th edition) to confirm probe placement for data inclusion^42^.

### Statistical analysis

Individuals conducting behavioral experiments were blinded to the pharmacological interventions. Results are expressed as means with standard error of the mean (M ± SEM), accompanied by individual data points for independent samples. Graphs were generated using GraphPad Prism 11 (Dotmatics, Boston, MA, USA). Statistical analyses were performed using SPSS v29.0.0.0 (IBM, Armonk, NY, USA). A paired t-test compared means between two related groups, and one-way ANOVAs compared means across three or more groups. The Holm-Sidak procedure controlled the familywise α level at 0.05 for multiple post hoc comparisons. Effect sizes were calculated using partial eta-squared (ηp²) for ANOVA main effects and interactions and Cohen’s d for key pairwise comparisons, including Student’s t-tests.

## Results

### Existence of a PACAP signaling network in the NAcc

We performed a series of experiments to establish the presence of the complete PACAP signaling mechanism in the NAcc. Figure 1 illustrates the molecular components of a PACAP signaling network detected in the rat NAcc. In Figure 1A, PAC1R protein expression is evident. The left lane in each image depicts the protein ladder corresponding to 50 kDa. A protein band labeled by the PAC1R antibody is visible at 53 kDa (Figure 1A, left image) but is absent in the presence of a PAC1R blocking peptide (Figure 1A, right image). In Figure 1B, the concentration of PACAP detected in microdialysis samples of the interstitial fluid of the rat NAcc is provided. Considering the possibility that PACAP can be transported across the blood-brain barrier at low concentrations^43,44^, we next confirmed its neuronal origin by assessing PACAP mRNA expression in NAcc afferents. We focused our measurements on the medial prefrontal cortex (mPFC) projections to the NAcc, as expression in cortical circuits may be relevant in comparison to the central sites of other gut-brain axis components (e.g., GLP-1). Figure 1C illustrates the experimental procedure involving the infusion of the retrograde tracer cholera toxin subunit B (CTb) into the NAcc. The localized spread of CTb was confirmed to the NAcc (Figure 1D). Retrogradely labeled cells expressing PACAP mRNA were detected in the prelimbic region of the mPFC (Figure 1E). The inset images show the co-expression of CTb and PACAP mRNA at a magnification of 40x. Collectively, these findings validate the existence of an endogenous central PACAP signaling network in the NAcc.

**Figure 1:**
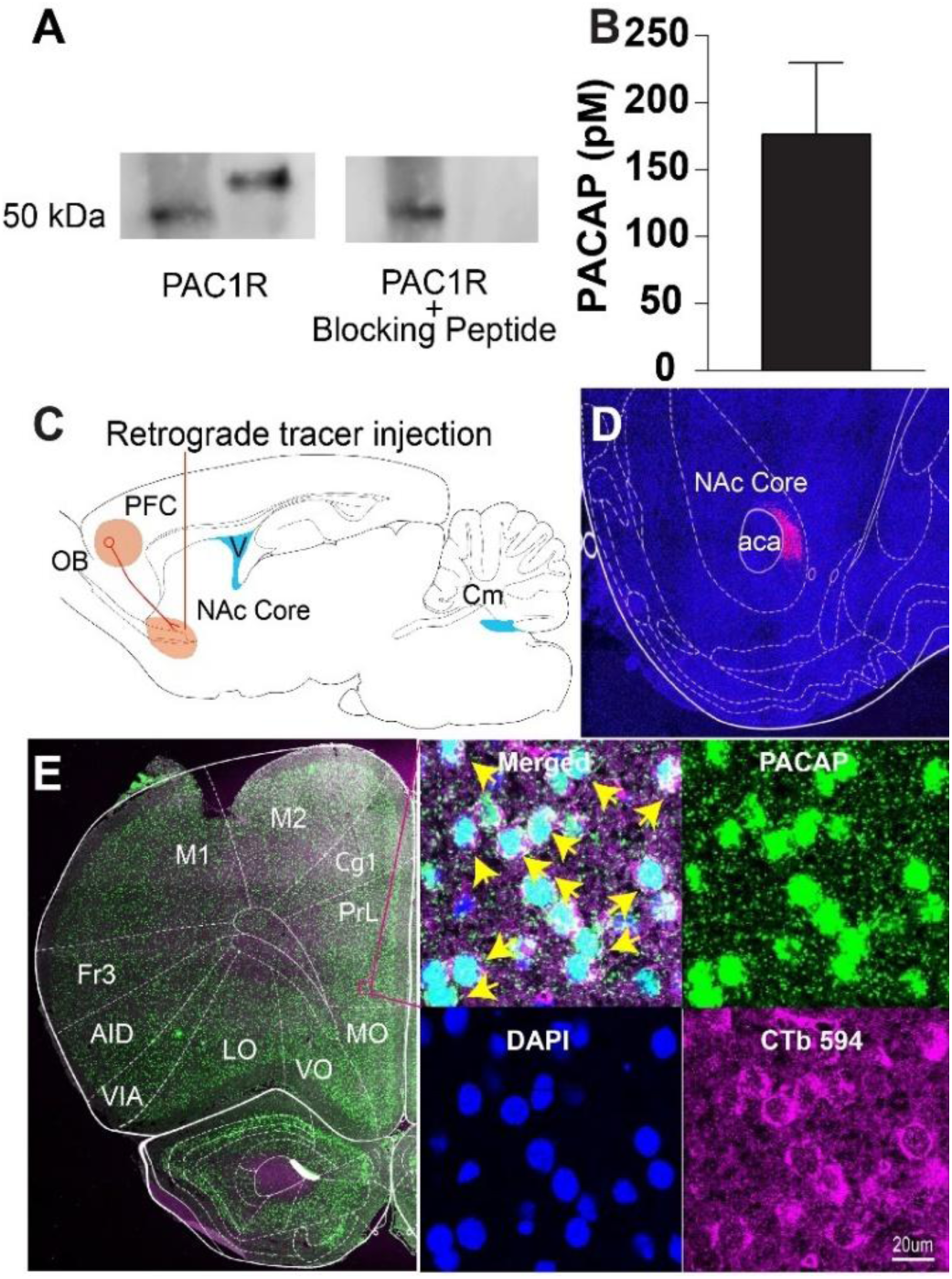
PACAP signaling in the NAcc. (A) Western blots of NAcc punch lysates confirming the presence of PACAP receptor (PAC1R), with specificity validated using a synthetic peptide. (B) Quantification of endogenous PACAP levels in the NAcc extracellular lysate by microdialysis (176.70 ± 53.27 pM, mean ± SEM). (C) Diagram illustrating retrograde tracing, highlighting the neural pathway from the prefrontal cortex (PFC) to the NAc core. (D) Confocal image (10×) of the NAc injection site showing the spread of cholera toxin subunit B (CTb) labeling (magenta) and cell nuclei stained with DAPI (blue), with the anterior commissure (aca) as an anatomical landmark. (E) Brain section highlighting prefrontal cortical regions with detailed anatomical labeling, including motor areas (M1, M2), orbital cortices (MO, VO, LO), and insular regions (AIV, AID). The inset (40× composite) shows PACAP mRNA expression (green), CTb-labeled projection neurons (magenta), nuclear DAPI staining (blue), and a merged image highlighting co-localization (yellow arrows). Abbreviations: OB (olfactory bulb), Cm (cerebellum), PrL (prelimbic cortex), Cg1 (cingulate cortex area 1), Fr3 (frontal cortex, area 3).

### PACAP microinjection in the NAcc does not stimulate cocaine-seeking behavior

In this experiment, we examined whether micro-infusions of PACAP (100 pM) are sufficient to reinstate extinguished cocaine-seeking behavior. Figure 2A depicts operant responding over a 2-hour period during the final extinction session (extinction) and during the test for reinstatement after an intra-NAcc infusion of PACAP (test). A paired sample t-test revealed no significant difference in lever pressing between the extinction and PACAP administration groups, t(30) = 0.749, p = 0.473 with a small effect size (Cohen’s d = −0.257). In Figure 2B, we depict the cannula placements, which were localized to the NAcc. These findings suggest that increased PACAP signaling in the NAcc does not promote drug-seeking behavior in rats with a history of cocaine self-administration.

**Figure 2:**
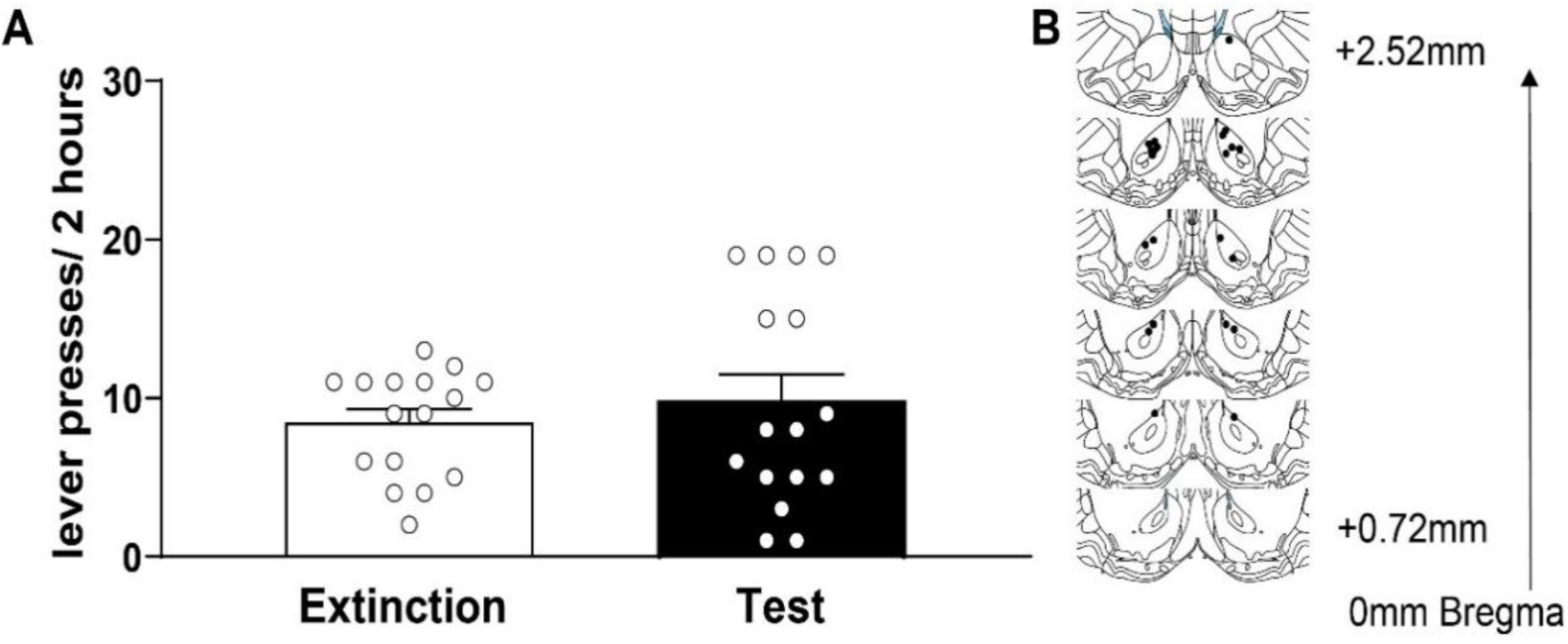
PACAP microinjection in the NAcc does not induce cocaine-seeking behavior. (A) Lever presses (mean ± SEM) by extinguished cocaine-seeking rats during a 2-hour session, recorded 10 minutes after PACAP (100 pM) microinjection into the NAcc. There was no significant difference in lever pressing between extinction and PACAP-treated groups, indicating that PACAP signaling in the NAcc does not promote cocaine-seeking behavior. (B) Schematic illustrating the PACAP injection sites within the NAcc, verified via histological analysis.

### Intra-PACAP infusion attenuates cocaine-primed reinstatement

To investigate the effect of intra-NAcc PACAP on cocaine-primed reinstatement, we administered vehicle or PACAP (50 or 100 pM) before reinstatement testing. A one-way ANOVA revealed a significant main effect of treatment (F_(3,27)_ = 11.161, p < 0.001; ηp² = 0.554), indicating a strong PACAP-mediated modulation of reinstatement. Post hoc pairwise comparisons confirmed a significant increase in reinstatement i.e., lever pressing following cocaine exposure (10 mg/kg) in the vehicle-treated group (Holm-Sidak, p=0.0001) demonstrating reinstatement of drug-seeking behavior. Rats infused with 50 pM PACAP (Holm-Sidak, p=0.0018) showed a partial attenuation of cocaine seeking. In contrast, administration of 100 pM PACAP blocked cocaine reinstatement since responding in this group was significantly lower than the vehicle + cocaine group (Holm-Sidak, p < 0.0008) and was not significantly different from extinction responding (Holm-Sidak, p>0.05).

Because the magnitude of reinstatement is sensitive to the amount of cocaine received during self-administration testing, we compared cumulative drug intake across the groups. A one-way ANOVA revealed no significant differences (p = 0.641), confirming comparable cocaine exposure across conditions (extinction: 1294 ± 76 mg/kg, vehicle: 1159 ± 66 mg/kg, 50 pM PACAP: 1210 ± 92 mg/kg, 100 pM PACAP: 1164.63 ± 125 mg/kg). Similarly, the number of extinction sessions required to meet extinction criteria did not differ among groups (p = 0.886; extinction: 8.90 ± 0.91, vehicle: 8.43 ± 1.78, 50 pM PACAP: 9.71 ± 1.36, 100 pM PACAP: 9.83 ± 1.72), ruling out extinction rate variability as a confounding factor. Cannula placement for PACAP microinjections were confirmed by Cresyl violet staining and only animals with accurate placement were used in the study (Figure 3B).

**Figure 3:**
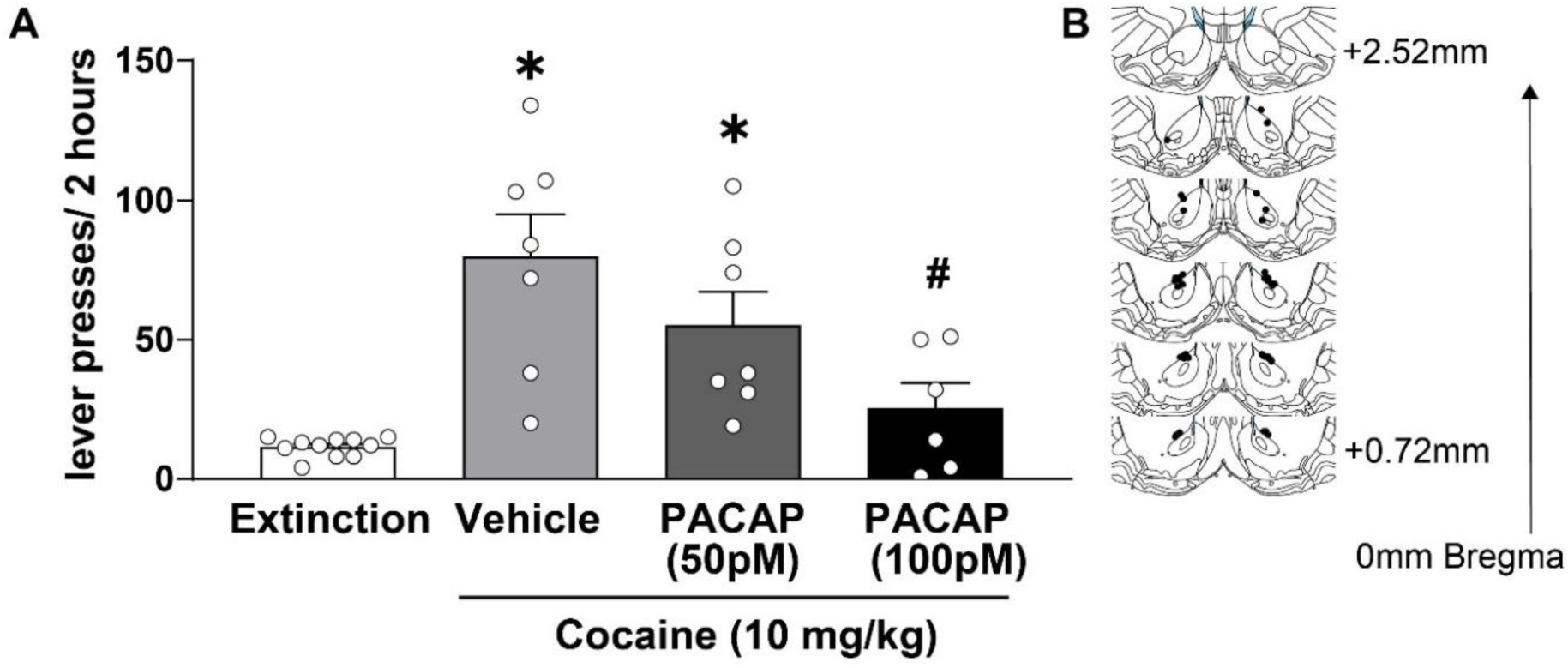
PACAP microinjection in the NAcc inhibits cocaine-primed reinstatement. (A) Lever presses (mean ± SEM) during a 2-hour session by extinguished cocaine-seeking rats reinstated with an intraperitoneal (i.p.) injection of 10 mg/kg cocaine. PACAP microinjected into the NAcc 10 minutes prior to the session significantly reduced cocaine-seeking behavior in a dose-dependent manner. Statistical significance was determined using the Holm-Sidak test. * and # indicate p < 0.05 compared to the extinction and vehicle groups, respectively. (B) Schematic showing the PACAP injection sites within the NAcc.

### PACAP attenuates D1 but not D2 receptor agonist-primed reinstatement

The NAcc integrates reward, executive function, deprivation states and other processes that regulate behavior^3,35–39^. One strategy to gain insight into which processes are being normalized by PACAP is to investigate the effect of NAcc PACAP signaling on drug seeking produced by intra-NAcc co-administration of a D1 or D2 dopamine receptor agonist, which have been shown to regulate distinct NAcc efferents linked to distinct processes regulating behavior^45–49^. Here we assessed the effects of intra-NAcc PACAP infusion co-administered with either the D1 receptor agonist SKF81297 or the D2 receptor agonist sumanirole maleate.

Figure 4A illustrates that intra-NAcc infusion of PACAP (100 pM) attenuated reinstatement primed by intra-NAcc infusion of the D1 receptor agonist SKF81297. A one-way ANOVA revealed a significant main effect of treatment (F_(2,30)_ = 18.21, p < 0.001,ηp² = 0.548), indicating strong PACAP-mediated modulation of D1-driven cocaine-seeking behavior. Post hoc pairwise comparisons confirmed that SKF81297 significantly increased reinstatement compared to extinction controls (p < 0.0001), demonstrating that D1 receptor activation enhances cocaine-seeking behavior. However, co-administration of 100 pM PACAP significantly attenuated SKF81297-induced reinstatement (p < 0.0003), suggesting that PACAP inhibits D1 receptor-mediated reinstatement. There were no significant differences in cocaine intake across groups (extinction: 1629 ± 84 mg/kg, SKF81297: 1676 ± 78 mg/kg, SKF81297 + PACAP: 1565 ± 85 mg/kg; p = 0.644). Similarly, the number of extinction sessions required to meet extinction criteria did not differ among groups (p = 0.104; extinction: 12.54±2.10, SKF81297: 9.08±2.42, SKF81297 + PACAP: 6.10±0.98). Cannula placement for PACAP microinjections were confirmed by Cresyl violet staining and only animals with accurate placement were used in the study (Figure 4B).

**Figure 4:**
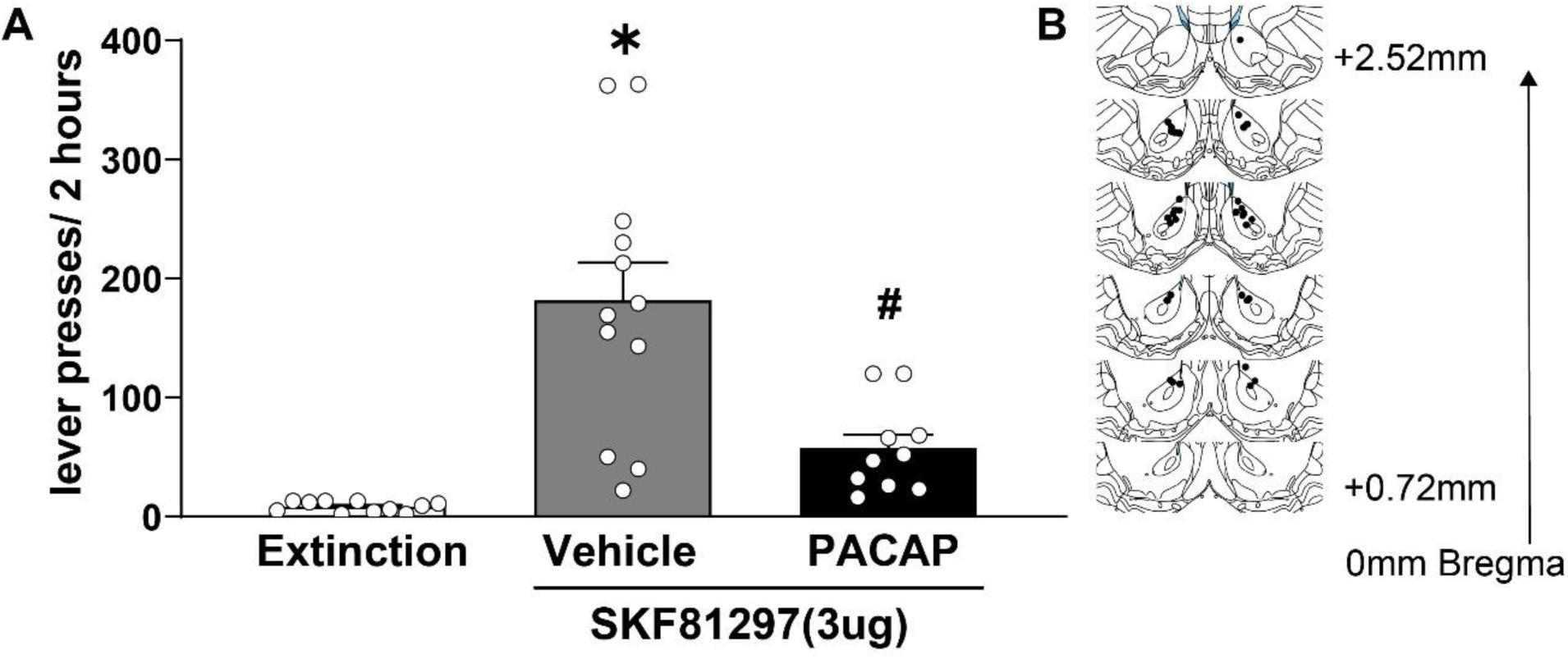
PACAP attenuates D1 receptor agonist-primed reinstatement. (A) Lever presses (mean ± SEM) during a 2-hour session by extinguished rats following intra-NAcc administration of the D1 receptor agonist SKF81297. SKF81297-induced reinstatement of cocaine-seeking behavior was significantly attenuated by PACAP (100 pM). Statistical significance was determined using the Holm-Sidak test. * and # indicate p < 0.05 compared to extinction and vehicle conditions, respectively. (B) Schematic showing the injection sites for SKF81297 and PACAP within the NAcc.

### D2 receptor agonist reinstatement

Next, we examined the effect of intra-NAcc PACAP infusion on reinstatement induced by the D2 receptor agonist sumanirole (Figure 5A). A one-way ANOVA revealed a significant main effect of treatment (F_(2,31)_ = 6.16, p = 0.006, ηp² = 0.284), indicating that D2 receptor activation produces a modest but significant increase in cocaine-seeking behavior. Post hoc Holm-Sidak pairwise comparisons confirmed that sumanirole significantly increased reinstatement compared to extinction controls (p = 0.0012), demonstrating that D2 receptor activation facilitates drug-seeking behavior. However, co-administration of 100 pM PACAP did not alter sumanirole-induced reinstatement, as lever pressing remained significantly elevated compared to extinction in the sumanirole + PACAP group (p = 0.0023) and did not differ from sumanirole alone (p > 0.05). These findings suggest that, unlike its effect on D1-mediated reinstatement, PACAP does not modulate D2 receptor-driven cocaine seeking under these conditions.

**Figure 5:**
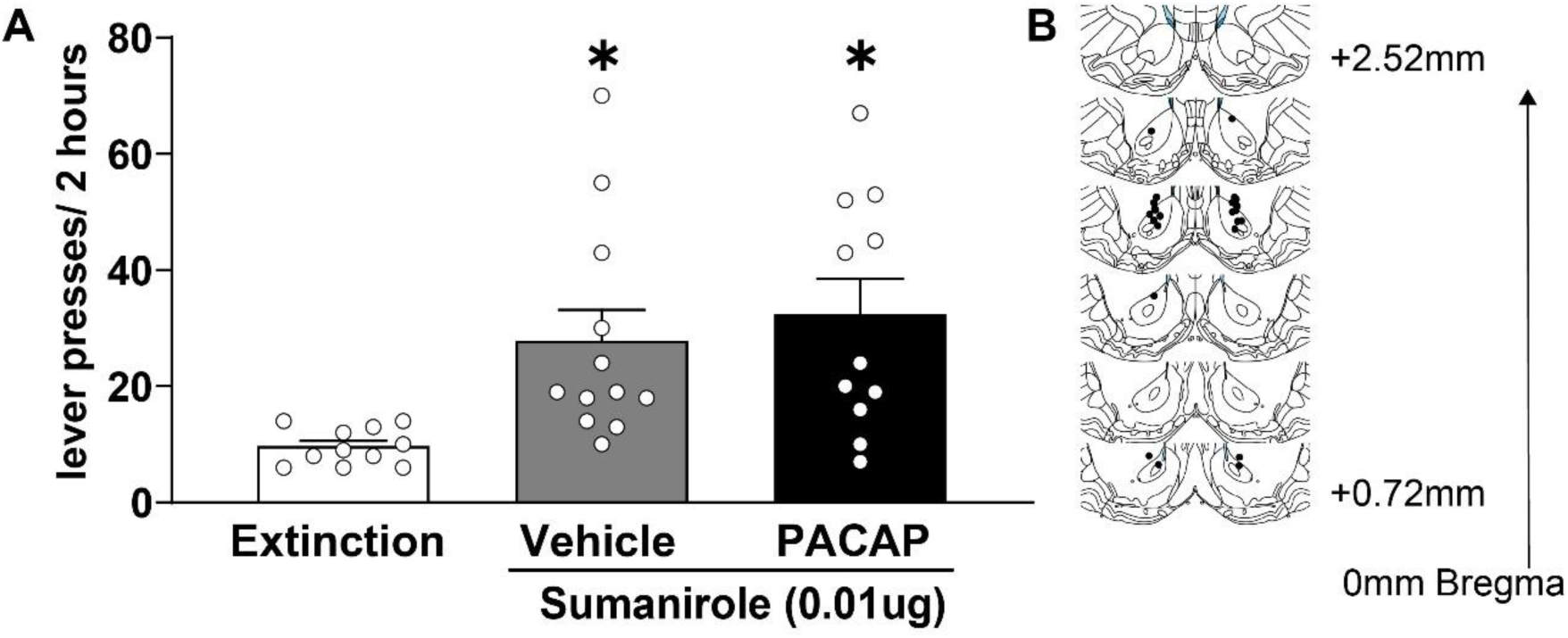
PACAP has no effect on D2 receptor agonist-primed reinstatement in drug seeking. **(A)** The data (mean ± SEM) show that administration of the D2 receptor agonist sumanirole moderately induces reinstatement in cocaine-experienced rats and PACAP (100 pM) co-administration does not inhibit it. Holm-Sidak test was used to determine significant differences between groups. * Indicates a p value<0.05 in comparison to extinction. **(B)** Schematic illustrates the sumanirole and PACAP injection sites.

To ensure that differences in reinstatement were not due to variations in prior cocaine exposure or extinction rates, we analyzed cumulative cocaine intake and extinction duration. A one-way ANOVA revealed no significant differences in cocaine intake across groups (p = 0.994; extinction: 1710 ± 60 mg/kg, sumanirole: 1702 ± 56 mg/kg, sumanirole + PACAP: 1710 ± 60 mg/kg). Similarly, the number of sessions required to reach extinction criteria did not differ across groups (p = 0.190), confirming that baseline drug intake and extinction learning were comparable across conditions. Cannula placement for PACAP microinjections were confirmed by Cresyl violet staining and only animals with accurate placement were used in the study (Figure 5B).

## Discussion

Our findings establish PACAP signaling as an integral component of the NAcc that selectively regulates drug-seeking behavior through dopamine receptor-specific mechanisms. Supporting this, we demonstrate that PACAP and its receptors are endogenously expressed in the rat NAcc core. Using retrograde tracing and in situ hybridization, we reveal the presence of PACAP mRNA-expressing neurons projecting from the PFC to the NAcc. In the cocaine self-administration/reinstatement paradigm, PACAP infusions into the NAcc did not reinstate cocaine-seeking in extinguished rats but blocked cocaine-primed reinstatement. Further, PACAP specifically inhibited reinstatement driven by intra-NAcc co-infusion of the D1 receptor agonist SKF81297, without altering behavior produced by intra-NAcc co-infusion of the D2 receptor agonist sumanirole. These results establish PACAP as an endogenous neuromodulator within the NAcc, selectively regulating motivated behavior by targeting specific dopaminergic pathways. By demonstrating PACAP’s ability to suppress cocaine-seeking through D1 receptor-dependent mechanisms and its localization in corticostriatal projections, our study highlights its potential role in CUD, a disorder characterized by dysregulated reward, emotional, and cognitive processes^1,2,4–9^.

Our first objective was to investigate the potential existence of a PACAP signaling network in the NAcc in rat. Western blot analysis confirmed the NAcc expression of PAC1R protein, a primary effector of PACAP signaling^50^. This is the first demonstration of PAC1R protein in the NAcc of Sprague Dawley rats, which extends existing observations documenting the presence of PAC1R protein in the NAcc of Wistar rats^33,34^. In vivo peptide microdialysis samples of NAcc interstitial fluid contained the PACAP peptide, marking the first direct evidence of extracellular PACAP in this region. Lastly, PACAP mRNA was detected in NAcc-projecting afferents from the medial prefrontal cortex, indicating that PACAP may be co-released with glutamate at corticostriatal synapses. This is an important finding because it suggests that neuronal release is a possible source of the extracellular PACAP detected in the NAcc. Alternative sources include transport across blood brain barriers, which can enter the rat brain after a bolus injection^51^. Future research is needed to examine the functional relevance of corticostriatal PACAP release to NAcc function. Collectively, these findings indicate the existence of a full signaling network within the NAcc.

The role of PACAP signaling in the NAcc on cocaine seeking behavior has not been investigated. Towards this, our findings reveal that intra-NAcc infusion of PACAP did not induce cocaine-seeking behavior but blocked cocaine-primed reinstatement (10 mg/kg, IP). These results align with prior work showing that PACAP signaling in the NAcc reduces hedonic drive^52,53^. Notably, the reduction of palatable food consumption by intra-NAcc infusion of PACAP mimicked the effect produced by increased GABA signaling in the NAcc^53^. While more work is needed, these findings suggest that PACAP may suppress synaptic transmission in the NAcc. Consistent with this, PACAP has been shown to increase the activity of astrocytic glutamatergic mechanisms, including GLT-1^54^ and system xc^55^, that decrease cocaine seeking^56–58^ likely through the suppression of synaptic transmission in the NAcc^59,60^.

Findings that may contrast with our results include the prior observations that reduced PACAP signaling through PAC1 receptor in the NAcc resulted in increased alcohol seeking. However, this outcome was strain specific as it is observed in Scr:sP but not Wistar rats^33,34^. Further, extrapolating from these findings to predict what occurs with increased PACAP signaling may be problematic. For example, the results from this study reflect the actions of PACAP at a single receptor, yet PACAP can signal through PAC1R and VIP receptors (VPAC1, VPAC2)^61^.

The brain wide effects of PACAP signaling on behaviors relevant for substance abuse reveal significant complexity. In the bed nucleus stria terminalis, PACAP signaling is sufficient to reinstate cocaine seeking and potentiate stress reinstatement of cocaine seeking^32^. Intracerebroventricular PACAP treatment accelerates morphine tolerance and enhances naloxone precipitated withdrawal^62^. The regional differences of PACAP on measures relative to substance abuse further support the need to identify manipulations that can target PACAP signaling in key regions, such as the NAcc, rather than those that broadly affect the entire brain.

The NAcc is a key region of interest as it integrates neural circuits from across the brain, playing a crucial role in cognition, emotion, reward, and other processes that shape behavior. One way to further explore the role of PACAP signaling in NAcc function is to examine how intra-PACAP infusions influence reinstatement induced by dopamine D1 or D2 receptor signaling in the NAcc as these receptors are believed to activate distinct circuits and functions^45^. Co-infusion of PACAP attenuated the robust reinstatement produced by intra-NAcc SKF81297 administration. In contrast, PACAP infusions did not alter the modest reinstatement produced by intra-NAcc infusion of sumanirole. While the precise functional distinctions between D1 and D2 signaling in the NAcc are still being investigated, it is evident that these receptors engage distinct processes underlying motivated behavior^45^. For example, Walle et al. (2024) found that selective chemogenetic activation of NAcc core D1-MSNs increased effort for a palatable reward but decreased overall food consumption, whereas D2-MSN activation increased immediate food intake while reducing effortful reward-seeking and locomotion^46^. D1 and D2 signaling within the NAcc and other striatal subregions also have distinct functions involving cognitive and reward processing^47–49^. Hence, these data indicate that PACAP in the NAcc acts in a precise manner by suppressing D1 receptor mediated processes without altering those mediated by D2 receptor activation.

Discrete components of the gut-brain axis appear to differentially target distinct mechanisms within the NAcc. While PACAP and GLP-1 belong to the same family of structurally related peptides that are distributed peripherally and centrally^24,25^, they may be producing complementary actions in the NAcc. Our findings demonstrate that PACAP signaling in the NAcc specifically regulates D1-mediated mechanisms to suppress cocaine seeking. In contrast, prior work implicates D2-expressing MSNs in behavioral control resulting from GLP-1 signaling in the NAcc. In support, neurons in the nucleus of the solitary tract (NTS) that produce GLP-1 innervate the NAcc^63^; GLP-1 signaling in the NAcc enhances glutamate release leading to activation of medium spiny neurons (MSNs) that underlies reduced feeding behavior^64^. Within the NAcc, suppression of behavior following increased MSN activity is most likely to involve MSNs expressing D2 dopamine receptors since suppression of MSNs expressing D1 dopamine receptors typically decreases motivated behavior^45,46,65,66^. Hence, PACAP, GLP-1, and potentially other components of the gut brain axis signaling in the NAcc may produce precise but complementary regulation of neural processes.

Our findings contribute to existing evidence establishing the strong therapeutic potential of the gut-brain axis as a therapeutic target for substance abuse. However, a deeper understanding of the peripheral and central mechanisms that facilitate these interactions is necessary to fully appreciate the potential of this new strategy. Our results emphasize the need to further investigate the interplay between PACAP and GLP-1 signaling in the NAcc and beyond, to determine whether they exert complementary or distinct influences on NAcc activity and behavior. Additionally, it is crucial to explore how peripheral mechanisms can selectively modulate PACAP and GLP-1 signaling within the NAcc. Unraveling these processes will be essential for fully evaluating the therapeutic potential of gut-derived interventions for cocaine use disorder and other conditions involving dysregulated behavioral control.

## Data availability statement

The data that support the findings of this study are available from the corresponding author upon reasonable request.

## Ethics approval statement

The experimental protocol involving animals were ethically reviewed and approved in accordance with guidelines for the care and use of laboratory animals, MU IACUC (AR-3815).

## Author contributions

D.B. conceptualized the study. D.B., C.S., H.E, K.L., and B.S. analyzed data and wrote the manuscript. B.S., S. G., H.E., K.L., M. B., R.N collected the data. All authors read and approved of the final manuscript.

